# Spinal lumen remodeling under the control of Gli signaling mechanically drives roof plate cells extension

**DOI:** 10.64898/2026.05.12.724326

**Authors:** A. Medyouf, A.M. Daza-Zapata, I. Anselme, A. Eschstruth, K.M. Kocha, P. Huang, S. Schneider-Maunoury, P.L. Bardet

**Affiliations:** Vertebrate brain morphogenesis team, Development, adaptation & aging (Dev2A) Laboratory, UMR8263, Sorbonne University, CNRS, Inserm, Paris, France; Paris Seine Biology Institute, Sorbonne University, CNRS, Inserm, Paris, France; Department of Biochemistry and Molecular Biology, Cumming School of Medicine, Alberta Children’s Hospital Research Institute, University of Calgary, Calgary, Canada

## Abstract

Morphogenesis often requires different cell types to coordinate their behaviors for an harmonious developpement. How these different cell type behaviors are synchronized within and between tissues remains one of the important questions to fully understand morphogenesis. We used the zebrafish developing spinal cord to study this question. At later stages of neurogenesis, the lumen of the neural tube remodels, by reducing its height dramatically to form the persisting ventral central canal, a morphogenetic process conserved in vertebrates. By combining genetics, cell signaling manipulation with antagonist drugs and high-resolution in vivo live imaging, we better characterised the dynamics and control of this remodeling process. We showed that the lumen retraction depends on Gli activity regulation, a downstream effector of the Shh morphogen signal. We further established that the lumen retraction is instrumental in the cellular elongation of spinal roof plate cells, a population that forms the ceiling of the spinal cord lumen. Our work therefore establishes that the Gli transcriptional regulators under the control of long-range morphogen Shh control lumen retraction and that this retraction is a key driver of the roof plate cells extension.

## INTRODUCTION

Morphogenesis is a tightly controlled process during metazoan development. The shape of growing tissues during development depends on a small set of cell behaviours that are controlled by cell-cell signaling pathways (Lecuit and Le Goff, 2007). Within a tissue, we start to better understand how biochemical and mechanical signals control precisely morphogenesis, like for example in lumen formation (Chan and Hiiragi, 2020). Yet, how two tissues made of different cell types that remodel within the same organ coordinate their morphogenesis is poorly understood and crucial to investigate.

To study the coordination of different cell types remodeling concomitantly, we focus on the vertebrate spinal neural tube. The neural tube is formed by radial glia, most of which are neuroepithelial progenitors that proliferate and differentiate to establish the entire diversity of cell types of the adult central nervous system (CNS) (Saade and Martí, 2025). The apical surfaces of radial glial cells surround a cavity also called lumen in the center of the neural tube, in which cerebrospinal fluid (CSF) circulates in the inflated part. Radial glia forming the neural tube of the CNS present primary cilia that bathe into the CSF (Kramer-Zucker et al., 2005). Cilia are essential in development ; they have a central role in orchestrating many sensory and signalling events of the cell (Andreu-Cervera et al., 2021; Bloodgood, 2009). In the CNS, primary cilia are essential for the differentiation and patterning of neural progenitors, whereas motile cilia are required for CSF circulation in brain ventricles and the spinal cord central canal (Kramer-Zucker et al., 2005; Ringers et al., 2019; Thouvenin et al., 2020). Cilia are therefore prime candidates to explore how cell signaling participates in coordination during vertebrate neural tube morphogenesis.

Shh is an important signaling pathway to control the proliferation and the patterning of differentiation of neural progenitors (Sagner and Briscoe, 2019). Shh morphogen is produced ventrally by floor plate cells and notochord and diffuses as gradient from ventral to dorsal in the neural tube (Andreu-Cervera et al., 2021). This Shh gradient is forming a Gli (glioma-associated oncogene) activity gradient along the dorso-ventral axis, that will regulate a series of target genes expression, including *patched-2* (*ptc2*). Gli transcriptional factors are the effectors of the signaling pathway, balancing between the Gli activator form (GliA) and the Gli repressor form (GliR) (Andreu-Cervera et al., 2021). The processing of these two forms requires the presence of a primary cilium. In zebrafish, mutants that lack primary cilia from early stage of development lose their Gli activity gradient, but maintain a residual low Gli1 activity - the activated form in zebrafish - that extends dorso-ventrally more than normal (Ben et al., 2011; Huang and Schier, 2009). Notch is another cell-cell signaling pathway that is crucial for neural progenitors to maintain both self-renewal and their apical domains forming most of the embryonic lumen border (Sharma et al., 2019). How these signals are involved in late morphogenesis events is not known.

In vertebrates, the CNS emerges from the embryonic neural tube, a pseudostratified epithelial structure with an internal lumen. At late stages of embryogenesis, the spinal cord central canal forms through a specific remodeling of the neural tube, namely the ventral retraction of the primitive lumen. This remodeling, sometimes called “dorsal collapse”, is conserved among vertebrates and has been observed in zebrafish, chicken, mouse, rat but also cat or camel (Böhme, 1988; Cañizares et al., 2020; Elmonem et al., 2007; Kondrychyn et al., 2013; Ribeiro et al., 2017; Ševc et al., 2009; Shinozuka and Takada, 2021; Sturrock, 1981; Tait et al., 2020). In the zebrafish, the primitive lumen extends over 80% of the total height of the neural tube, although only the most ventral part is inflated by the circulating CSF (Figure S1C) (Thouvenin et al., 2020). Between 2 and 3 days of development, this primitive lumen shrinks to the ventral part of the neural tube to form the central canal (Figure 1 A,B) (Ribeiro et al., 2017). Lumen remodeling is the scene of various concomitant cellular rearrangements : dorsal lateral progenitors (also called dVL for “dorsal ventricular layer”) stop contributing to the formation of the lumen border by detaching their apical surfaces (a process sometimes called “cell attrition”), while the roof plate cells that form the dorsal midline of the neural tube (also called dmRG, for dorso-medial radial glia), adapt to this reduction and elongate ventrally to several times their initial size (Böhme, 1988; Kondrychyn et al., 2013; Shinozuka et al., 2019; Tait et al., 2020). The lateral progenitors and dorsal roof plate cells thus remodel in a coordinated manner during this period. Tait et al. (2020) proposed a model, in mouse and chicken, where roof plate cells promote the loss of apical polarity of the most dorsal lateral progenitors forming the lumen by secreting CRB2S molecules, leading to their delamination and the reduction of lumen height.

**Figure 1:**
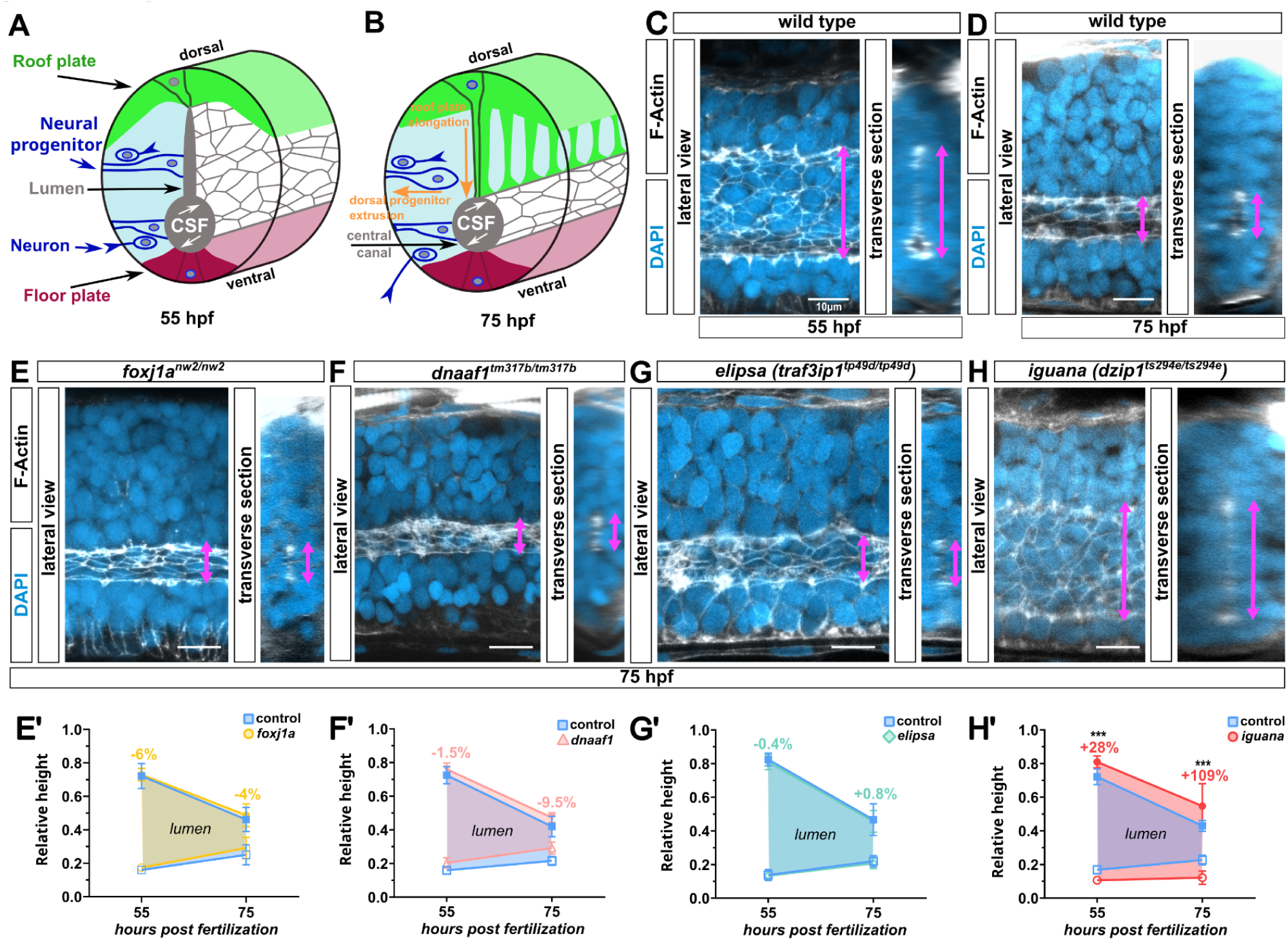
Lumen remodeling is not impaired in the absence of cilia, but is altered in iguana mutants. **A.** 3-D diagram of the neural tube at 55hpf depicting the full extent of the spinal lumen formed by the lateral progenitors apical surfaces (blue). It is connected dorsally to the roof plate (green) and ventrally to the floor plate (purple). **B.** 3-D diagram of the neural tube at 75hpf depicting the spinal central canal after remodelling. Orange arrows represent the typical cell behaviours associated with lumen height reduction : progenitor extrusion and roof plate elongation. **C-H.** Confocal images of F-actin delineating the neural progenitors apical surfaces forming the lumen (phalloidin staining, white) and of the neural tube nuclei (DAPI in blue) in zebrafish embryonic spinal cords at different hours post-fertilisation (hpf). The pink arrows indicate the dorso-ventral extent of the lumen. **C.** Neural tube at 55hpf, the lumen is still extended dorsally. **D.** Neural tube at 75hpf, lumen has been remodelled to form the central canal in the ventral part of the neural tube. **E, F, G, H** Confocal images of a 75hpf spinal neural tube of foxj1a (E), dnaaf1 (F), elipsa (G) and iguana (H) mutants. **E’, F’, G’, H’.** Graphical representation of the relative lumen height at 55hpf and 75hpf comparing control siblings (blue) to foxj1a homozygous mutants (E’, yellow, n=14 at 55hpf, n=19 at 75hpf), dnaaf1 (F’, pink, n=14 at 55hpf, n=21 at 75hpf), elipsa (G’, green, n=10 at 55hpf, n=19 at 75hpf) and iguana (H’, red, n=13 at 55hpf, n=32 at 75hpf). Relative lumen height is highlighted with a coloured area, while the floor plate and roof plate relative heights are below and above coloured area respectively. Percentages represent the relative lumen heights difference between the mutant and the control. *** = p < 0.0001.

To improve our understanding of behaviour coordination of various cell types within an organ, we decided to use the zebrafish embryo spinal cord remodeling as a model to perform live confocal imaging and functional studies. We sought to better understand how cell-cell signaling regulates the spinal lumen retraction and its coordination with roof plate cells elongation.

We first turned our attention to cilia as an important relay in multiple signaling pathways. By studying different zebrafish mutants defective in cilia functions, we demonstrated that cilia absence does not directly impact lumen remodeling but that *dzip1* mutant embryos (*iguana*) - encoding the Dzip1 protein located at the ciliary basal body - present a defect of lumen remodelling. While characterising the *iguana* mutant phenotype, we observed an expansion of *ptc2* expression, a well known target of the Gli activity in this mutant. Our findings demonstrate that Gli activity needs to be shut down for dorsal progenitors to withdraw from the lumen border, hence leading to lumen retraction. Indeed, modulation of Gli activity in embryos impacted lumen remodeling. Over-expression of the Gli activator reproduced the *iguana* defect of lumen retraction while over-expression of Gli repressor and inhibition of Gli activator rescued the *iguana* phenotype. By artificially accelerating progenitor extrusion via blocking Notch signaling, we uncover a mechanical coupling between the retracting lumen border and roof plate cell extension. We therefore demonstrate that progenitor cells extrusion from the lumen border under Gli control is the driver of lumen retraction and, via mechanical coupling, of roof plate cell extension.

## MATERIALS AND METHODS

### Zebrafish strains

All zebrafish strains used in this study were maintained and raised under standard conditions. Animals were raised at 28.5 °C under a 14/10 light/dark cycle until the start of the experiment. Eggs were collected an hour after mating and maintained at 28.5 °C in E3 medium (2011 Cold Spring Harbor Laboratory Press). All procedures were performed on zebrafish embryos between 30hpf and 95hpf. All experiments were performed on Danio rerio embryos of AB, Tüpfel long fin (TL) and nacre background. The transgenic strains used were: *TgKI(tjp1a-eGFP)^pd1252^*, *TgKi(tjp1a-tdTomato)^pd1224^* (Levic et al., 2021), *Et(krt4:EGFP)^sqet33^* (Kondrychyn et al., 2013), *Tg(hsp70l:GFP-gli1)^a4594^*(Huang and Schier, 2009), *TgBAC(ptc2:Kaede)^a4596^* (Huang et al., 2012), *Tg(hsp:Gli3R-EGFP)^ca121^* (see below). The *traf3ip1^tp49d/tp49d^*, referred to as *elipsa* mutant (Driever et al., 1996), foxj1a^nw2/nw2^, referred to as *foxj1a* mutant (Thouvenin et al., 2020), *dnaaf1^tm317b/tm317b^*, referred to as *dnaaf* mutant (Brand et al., 1996; Van Rooijen et al., 2008), and *dzip1^ts294e/ts294e^*, referred as *iguana* mutant (Brand et al., 1996) mutant strains were maintained as heterozygotes, and homozygous embryos were generated by intercrossing heterozygous carriers.

### Building of the *Tg(hsp:Gli3R-EGFP)^ca121^* transgenic line

To generate the hsp:Gli3R-EGFP construct, a C-terminally truncated zebrafish gli3 gene fused to EGFP (Huang and Schier, 2009) was cloned downstream of the hsp70 promoter. To establish a stable transgenic line, 40 pg of hsp:Gli3R-EGFP plasmid DNA was co-injected with 40 pg of tol2 transposase mRNA into wild-type embryos at the one-cell stage. Positive transgenic lines were identified by screening for EGFP expression in F1 embryos following heat shock.

### Genotyping

Homozygous ciliary mutant embryos for *elipsa, dnaaf1, foxj1a* and *iguana* were identified by observation of curled down embryos after crossing, leading to identify adult heterozygous zebrafish. For *dnaaf1* and *foxj1a*, primers were also designed to identify the mutated loci in adult zebrafish. DNA of anesthetised adult fish were extracted using fin clipping. The fin piece was then digested for 3 hours at 55 °C in 100ul of fin clip medium (100mM de Tris-Hcl pH7,5, 1mM EDTA, 250mM NaCl, 0,2%SDS) + 1ul of PKA (0,1µg/µl). Amplification of the genomic DNA were performed using the following primers for *foxj1a*, forward-mutation 5’ -TCTCCATCCTCAACGCCAGG- 3’ and reverse 5’ -ACTTCGTCGGGACAGTGTCC-3’ and *dnaaf1* forward 5’ -GCAAGCTTTGCACGCTTAATGTCTC- 3’ reverse-mutation 5’-GTCACAAACATTCTCCAGTGTT- 3’. For hsp:Gli3R-EGFP and hsp:EGFP-Gli1 transgenic lines were genotyped by eGFP PCR detection using the following primers : forward 5’-GCATGGACGAGCTGTACAAG- 3’ and reverse 5’ -TGAACAGCTCCTCGCCCTTG- 3’

### Immunostaining

Embryos were manually dechorionated and anesthetized using 0.4% tricaine (MS 222, E10521, Sigma) for 2 minutes prior to fixation. All the following steps are done with agitation. Embryos were then fixed over-night in 4% PFA 1,5% saccharose or 4h at room temperature (RT), rinsed 5 times for 15 mins in 1X PBS. Embryos were blocked at least 1h at room temperature in a solution containing bovine serum albumin (BSA, 2mg/ml), 0.7% Triton, 1% DMSO, and 10% normal goat serum (NGS). Embryos were incubated with primary antibodies 48h at 4 °C in a solution containing BSA (2mg/mL), 0.5% Triton, 1% DMSO, and 1% NGS and subsequently washed four times during one hour in a PBS 1X 0.5% Triton. Secondary antibodies were added overnight at 4 °C in a solution containing 0.5% Triton, 1% DMSO, and 1% NGS. Embryos were then washed 4 times for 15mins using 1X PBS 0.5% Triton. The yolk and the head of the zebrafish embryo were removed, and the trunk was mounted laterally in Vectashield Antifade Mounting Medium (Clinisciences, H1000). The following dilutions of primary antibodies were used: chicken IgY anti-GFP 1:250 (AVES Lab), mouse IgG1k anti-ZO-1 1:250 (Zymed-Invitrogen), mouse IgG2b anti-Acetylated-tubulin 1:300 (Sigma), mouse IgG1k anti-GT335 (AdipoGen LifeSciences), rabbit polyclonal anti-Sox2 (GeneTex), mouse monoclonal anti-HuC/HuD (Invitrogen LifeTechnologies), mouse IgG1 anti-Nkx6.1 1:150 (DSHB), phalloidin alexa 568 1:400 (Invitrogen) and DAPI 1:1000. All secondary antibodies were purchased from Molecular Probes and The Jackson Laboratory and used at 1:200.

For HuC staining, to improve antibodies penetration, embryos were cut in two at the cloaca level after PFA fixation and incubated 7mins in acetone at -20°C before following the usual protocol.

### Image acquisition

Fixed embryos were imaged on a Zeiss LSM 980 Upright Confocal microscope with 63X Oil magnification. The images were all acquired at a fixed rostro-caudal position, at the level of the cloaca in the dorsal part. We acquired Z-stacks of the neural tube to obtain a 3D stack acquisition in a lateral view, except for HuC stainings where the embryos were mounted head down in agarose to acquire transverse sections. Images were then processed using Fiji.

For live imaging, embryos at the appropriate stage were manually dechorionated, anaesthetised with 0.4% tricaine and then embedded in 0.5% low melting agarose in 1X E3 medium on 1% Agarose E3 mould. Movies were recorded on a Zeiss Spinning Disk upright confocal microscope using a 40X water lens, one image every 15 minutes were taken.

### Quantification of the different neural tube heights

All images were treated with Fiji software. We used digital reslicing to reconstruct a series of transverse sections of the neural tube. We determined manually the height of all regions composing the neural tube (neural tube height, roof plate length, lumen height, floor plate length) using Fiji straight line measure, using usually at least 3 sections per embryo that represents internal diversity. The limit of each domain was determined by the intercept between the dorso-ventral axis and a perpendicular line at the level of the extremity of the signal delineating the neural tube and the lumen. For Sox2 and Nkx6.1 stainings, the same was done with a perpendicular line marking the limit of the expression domain. Prism software (GraphPad) or Matlab (MathWorks) were then used to compute the heights of the different regions of interest using the bottom of the neural tube as a reference at different stages (hpf) and condition (mutant vs control), plot them and compute statistical comparisons.

### Specificities for quantification of live imaging movies

After measuring the different neural tube heights on images from live time-lapse movies, we used Matlab to plot them and measure the evolution of lumen height, and fit a linear curve to calculate the speed of lumen retraction. In the case of the *iguana* movies, we noticed that the control embryo had a lower speed of retraction when compared to a fixed sibling raised at 28°5C. This was due to a colder temperature (around 24°C) in the microscope room. We therefore corrected the time step of this experiment so the live control embryos fit the fixed controls. This correction was confirmed by the fact that after this correction, the *iguana* homozygote mutants imaged in the same dish had the same speed of lumen retraction as fixed embryos raised at 28°5C.

### Cilia density counting and normalisation

Acetylated tubulin and GT-335 antibodies were used to stain cilia. Cilia density was calculated by counting the number of cilia in 100x100 pixels square in each image. All data were normalized by the means of controls.

### Segmentation

To quantify the area and elongation of the apical surface of the lumen cells we used a combination of MATLAB software algorithm (The MathWorks) and Fiji analysis. Due to technical limitations, only the most ventral cells, at the level of the inflation of the lumen, were amenable to a Z-projection compatible with a segmentation. First we created one stack image of the apical surface of the progenitors (using either max or average projection) on Fiji. These images were then segmented using MATLAB script (Bardet et al., 2013). Finally quantification of the area and cells elongation of each cell were performed thanks to MorphoLib plugin (https://github.com/ijpb/MorphoLibJ)

### Counting apical surface of progenitors forming the lumen

To determine the number of progenitor apical surfaces contributing to the lumen on one side of the border along the dorso-ventral axis, we used Z-stack of ZO-1 or phalloidin staining on fixed embryos at different stages. The delineation of the apical junction belt by ZO-1 or phalloidin allowed us to determine the outline of each progenitor apical surface, while moving in 3D through the Z-stack. We counted 3 positions per embryo along the rostro-caudal axis of the neural tube. The mean of these three counts is the mean of the number of apical surfaces forming the lumen.

### Cerebrospinal fluid (CSF) labelling

49hpf, 54hpf, 74hpf embryos were manually dechorionated, mounted in 0.5% low-melting point agarose in the lateral position covered by E3 medium 0.4% tricaine to anaesthetize the embryos. A few nanoliters containing Rhodamine Dextran 10 000 MW (D1817, Invitrogen) at 5 mg/ml diluted in E3 medium were injected in the pericardiac chamber in order to label CSF. Following the injection, we waited one hour before performing experiments to allow the particles to be transported down the central canal. We acquire live embryos to access the height of the circulating flow in control vs mutant embryos.

### PHotoconvertile REporter of Signalling History (PHRESH) analysis

All fluorescent imaging was carried out using the Zeiss LSM980 Upright confocal microscope and the Zeiss software. Photoconversion was carried out using the 405 nm laser with a 20x water-dipping objective. ptc2:Kaede embryos at the appropriate stages were anaesthetised with 0.4% tricaine and then embedded in 0.5% low melting agarose. To achieve complete conversion over a large area, a rectangular area of 1000 by 300 pixels was converted by scanning the area with 50% 405 nm laser at 200 ms per pixel. Following confirmation of Kaede-red expression by acquisition, embryos were recovered in E3 medium for 20hours post-conversion before imaging again the same region. Appropriate imaging parameters were established using the unconverted region as a reference to avoid over or under exposure of the Kaede-green signal. Cross-sections were generated using Fiji-ImageJ software (Schindelin et al., 2012) to create a 3D reconstruction of the image, then ‘resliced’ to yield transverse views of the spinal cord.

To generate PHRESH signalling profiles from the reconstructed transverse views along the dorso-ventral axis, three boxes were drawn directly through average projections of transverse sections stacks of the spinal canal and the fluorescent intensity of Kaede-green was measured along long axis of the box. Measurements were taken at three different positions of the neural tube, and the average of the three boxes was presented in the graph. Distance in µm were normalised per the neural tube height to allow comparison between fish with slight variations of the total neural tube height, and interpolation used to replot the profile with the same distance step for all embryos. These profiles were generated using Fiji-ImageJ software then graphically represented using GraphPad.

### Drug Treatment

Embryos at the appropriate stage were treated with LY-411575 (Sigma, 25µM), GANT-61 (Sigma, 80µM) or DMSO in E3 medium. For the live LY-411575 experiment, the drug was added 30 minutes before the beginning of acquisition. For the fixed experiment, the drug-treated larvae were kept in the dark at 28°C until the time of fixation.

### Heat Shock experiments

To induce expression from the heat shock promoter, embryos at the relevant stage were placed in a 2 ml tube with 1ml of E3 medium in a water bath set to 37,5 °C for 1hr. For hsp:EGFP-Gli1 experiment, embryos were placed at 28°C from 0hpf to 30hpf, then at 25°C from 30hpf for 18hours to have embryos at 40hpf stage at 9am. Two heat shocks were then performed, one at 40hpf stage and the other at 50hpf stage. Embryos were transferred back into E3 medium in a petri dish and recovered at 28 °C until 70hpf stage for fixation. Between the two heat shocks, embryos were placed back to E3 medium at 28°C. For hsp:Gli3R-EGFP, one heat shock was performed at 50hpf. After heat shock, embryos were transferred back into E3 medium in a petri dish and recovered at 28 °C until 70hpf for fixation. For drug treatment after heat shock, embryos were transferred directly from the heat shock to E3 medium containing the appropriate drug.

### Statistics

Statistical details related to sample size and p values, are reported in the figure legends. All statistics were performed using Prism or Matlab. In the figure panels, asterisks denote the statistical significance calculated using the appropriate test (stated for each test in the legends): * = p<0.05; ** = p<0.01; *** = p<0.001; **** = p<0.0001; ns = p>0.05.

## RESULTS

### 1. Cilia are dispensable during lumen remodeling, but Dzip1/*iguana* function is required

We first explored whether cilia were involved in cell signaling required to control spinal remodeling coordination. We noticed that the inflated ventral part of the lumen seems to coincide with the final border of the central canal, and we thought that circulating CSF ventrally might be a cue to determine the upper limit for lumen border retraction (Figure 1A-D, Figure S1C compare 55 hours post fertilization (hpf) and 75hpf, (Ribeiro et al., 2017). We therefore focused on mutants affecting cilia motility only, since they do not properly inflate the ventral circulating portion of the lumen early on (Thouvenin et al., 2020). We analysed two mutants defective for cilia motility : *foxj1a* (*foxj1a^nw2/nw2^*) (Thouvenin et al., 2020), which affects a master transcription factor driving cilia motility, and *dnaaf1,* (dnaaf1^tm317b/tm317b^) (Brand et al., 1996) that encodes dynein axonemal assembly factor 1, involved in cilia motility. We did not observe defects of lumen border height during (55hpf) and at the end (75hpf) of lumen remodeling in the *foxj1a* and *dnaaf1* mutants (Figure 1E-F and 1E’-F’ for quantification). This indicates that CSF circulation and motile cilia are dispensable for proper lumen border remodeling in the zebrafish spinal cord.

We then focused our attention on mutants with defects in primary cilia, known to be important relays in different signaling pathways during neurogenesis (Andreu-Cervera et al., 2021). We studied the *elipsa* mutant (*traf^3ip1^*) (Driever et al., 1996; Bizet et al., 2015) and the *iguana* mutant (*dzip1^ts294e^*) (Kim et al., 2010). In the *elipsa* mutant embryo, that affects a protein involved in the intraflagellar transport (IFT) and leads to loss of both motile and primary cilia (Omori et al., 2008), we observed a normal lumen height at the two timepoints we analysed (Figure 1G, 1G’). As this mutant lacks all cilia at 55hpf - the onset of lumen remodeling (Omori et al., 2008) (Figure S1A-B), this indicates that cilia are dispensable at the onset of and during lumen retraction.

Contrary to the other mutants we studied, *iguana* mutant embryos presented a defect in lumen remodeling. The *iguana* mutant (*dzip1^ts294e^*) is defective for the Dzip1 protein that localises to the basal body of cilium, causing defects in primary ciliogenesis (Kim et al., 2010). At 55hpf, the lumen border of *iguana* mutants was already 28% higher than that of control siblings (Figure 1H-H’). This difference was further increased at the end of remodeling at 75hpf, as the lumen border was on average two times higher in *iguana* embryos than in controls with a higher variability of lumen height in iguana embryos (Figure 1H-H’, see also Figure S2A, dispersion at 75hpf). We therefore concluded that Dzip1 is required for spinal lumen remodeling. Contrary to *elipsa*, the *iguana* mutant loses most of its primary cilia at an early stage, and exhibits a change in Gli activity regulation that other ciliary mutants don’t have (30hpf - Figure S1A-B) (Huang and Schier, 2009; Kim et al., 2010; Omori et al., 2008).

Moreover, previous works showed the role of motile cilia in central canal inflation (Thouvenin et al., 2020). We therefore compared central canal inflation in *iguana and elipsa* mutants. We observed a decrease in the diameter of the inflated part of the canal in both ciliary mutants at all stages, further demonstrating that canal inflation has no impact on lumen remodeling and retraction (Figure S1C-E). Altogether, our work points towards a specific requirement for the *dzip1* gene in lumen border remodeling.

### 2. Lumen remodeling in *iguana* presents changes in apical surface shape and behavior of progenitors

To better understand the lumen border height difference between *iguana* and control sibling embryos, we looked at the dynamics of lumen remodelling. We used a double transgenic line expressing GFP specifically in roof plate cells cytoplasm (Et(krt4:eGFP)^sqEt33^, Kondrychyn et al., 2013), together with a fusion of the tight junction protein ZO-1 with tdTomato, to outline the apical surfaces of the neural progenitors forming the lumen border (*TgKi(tjp1a-tdTomato)^pd1224^*, Levic et al., 2021). We followed lumen border remodeling by real-time confocal imaging. Control siblings exhibited a regular lumen border retraction between 55hpf and 75hpf (Figure 2A, blue curve in B for quantification ; average retraction speed close to 1 µm/h, see Movie S1 and Figure S2A-B). Spinal cords of *iguana* mutants displayed normal proportions of lumen border height at the beginning of our real-time experiments (50hpf, Figure 2A’-B and Figure S2B-D). However, the retraction of the lumen border was slower (Figure 2A’, red curve in B for quantification ; average retraction speed around 0.5µm/h, see Figure S2B). These observations are consistent with our time course experiments on fixed embryos (from 35hpf to 95hpf, Figure S2A-C) (Ribeiro et al., 2017). Despite a slightly shorter neural tube in *iguana* mutants (Figure S1E’), the lumen border absolute height and its proportion of the neural tube were significantly higher in *iguana* compared to wild-type siblings from 55hpf onward (see quantifications on Figure S2C). The lumen border of *iguana* mutants never retracted to the proportion observed in wild-type until as late as 95hpf, when remodeling was long finished in the control siblings (Figure S2A-D). Note that the progression of the roof plate extension was also slower in the *iguana* mutants compared to control siblings (Figure 2A-A’, 2B), supporting a coupling between the remodeling of these tissues (see below, Figure 4). These results confirm that Dzip1 is required for proper control of lumen border retraction.

**Figure 2.**
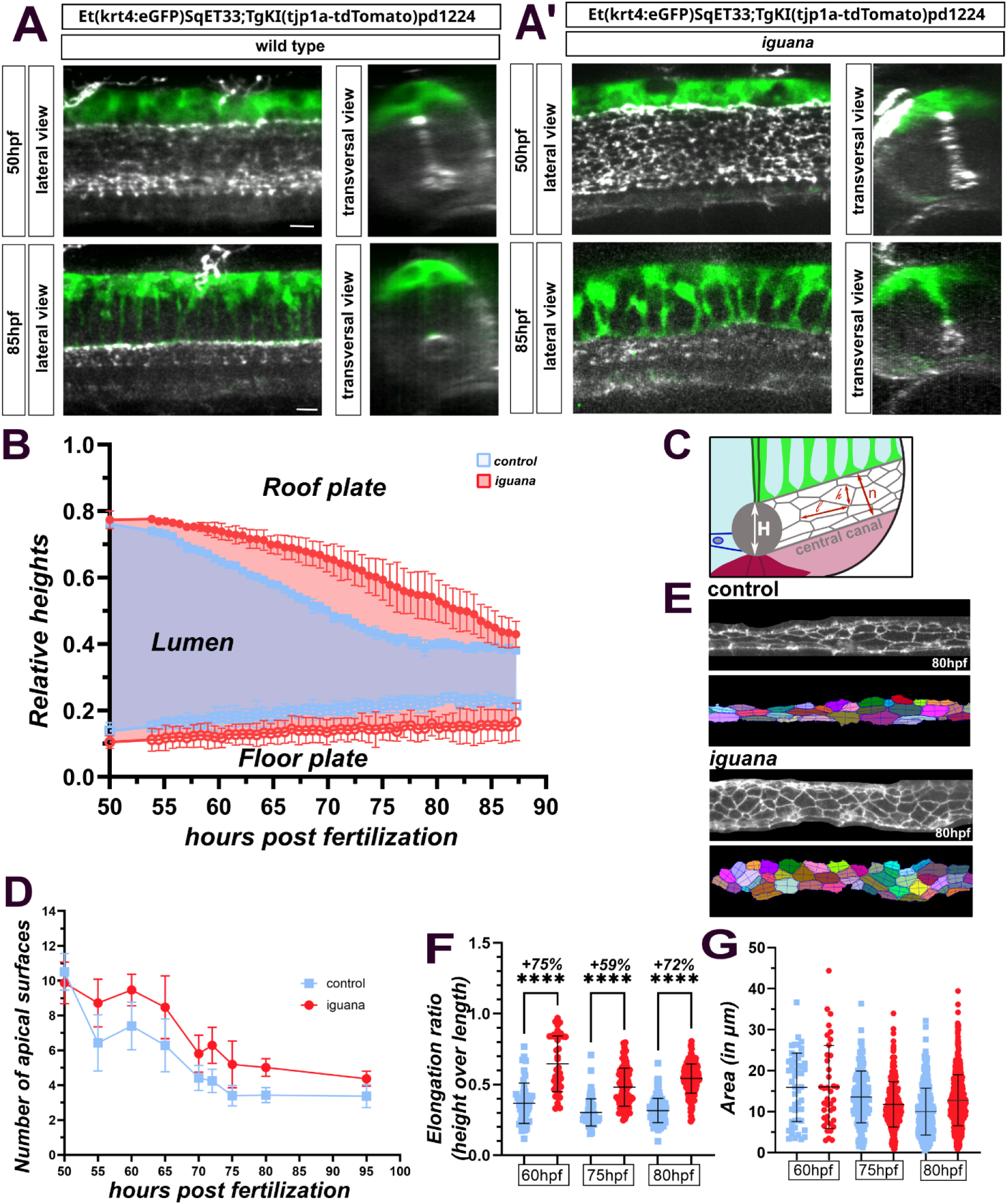
Iguana defect is due to an increased number of progenitors contributing to the lumen together with a change in their apical surface shape. **A-A’.** Lateral and transversal views of live Et(krt4:eGFP)^SqET33^;TgKI(tjp1a-tdTomato)^pd1224^ embryos at 50hpf and 85hpf of control (A) and iguana mutant (A’). **B.** Quantification of lumen remodelling live in control (blue, n=2) and iguana embryos (red, n=8) from 50hpf to 87hpf. **C.** Scheme of central canal, presenting the height of the lumen, H, dependent on the number of progenitors composing the lumen, n, and the height of their apical surface, h. **D.** Number of progenitor apical surfaces, outlined by ZO-1 staining, composing the lumen during remodeling from 50hpf to 95hpf in control (blue, n=13 at 50hpf, n=11 at 55hpf, n=8 at 60hpf, n=14 at 65hpf, n=6 at 70hpf, n=15 at 72hpf, n=16 at 75hpf, n=4 at 80hpf, n=4 at 95hpf) and iguana (red, n=9 at 50hpf and 55hpf, n=12 at 60hpf, n=8 at 65hpf, n=13 at 70hpf, n=18 at 72hpf, n=23 at 75hpf, n=5 at 80hpf, n=4 at 95hpf). **E.** Example of segmentation of apical surfaces of lumen-forming progenitors at 80hpf in control and iguana. **F.** Elongation ratio (height/length) of the progenitors apical surfaces and **G.** Area in µm of the progenitors apical surfaces forming the lumen in iguana (red) and control siblings (blue) at 60hpf (n=1 for control and iguana), 75hpf (n=1 for control and n=3 for iguana) and 80hpf (n=4 for control and iguana).

As the spinal lumen border is composed of the apical surfaces of neural progenitors, we reasoned that a higher border in *iguana* could be the consequence of an increased number of apical surfaces or of higher apical surfaces, or both (Figure 2C). We counted the number of progenitor apical surfaces contributing to the lumen border from 50hpf to 95hpf in both *iguana* embryos and control siblings (Figure 2D). While there was no difference at 50hpf, before the onset of lumen retraction, *iguana* mutant lumen borders rapidly displayed an excess of 1 to 2 progenitor apical surfaces forming the lumen border (Figure 2D). We also noticed a larger of Sox2+ domain along the spinal lumen from 60hpf onward (Figure S2E-H), suggesting that the *iguana* mutant has indeed an excess of polarised neural progenitors persisting in time. We then evaluated the shape of the apical surfaces of the most ventral progenitors, that eventually persist to form the central canal (Figure 2E). From 60hpf to 80hpf, apical surfaces of *iguana* mutants consistently displayed a greater height/length ratio (Figure 2F), meaning they were less elongated along the rosto-caudal axis but the apical surface did not present any change in their area quantification (Figure 2G).

The defects in lumen border height in *iguana* mutants can therefore be explained by an increased number of progenitor apical surfaces forming the lumen and by a greater height-to-length ratio of these surfaces (Figure 2D, 2F, 2G). Dzip1 is hence required to regulate both the shape of apical surfaces and the number of progenitors contributing to the lumen border.

### 3. Gli activity at the onset of lumen remodeling controls the final height of the central canal

As shown previously (Figure 1), we have excluded a direct requirement for cilia during the lumen border remodeling process itself. However, previous observations in *iguana* mutants reported a dorsal shift of medium-range Shh targets like Olig2 (Jacobs and Huang, 2019) that is thought to depend on Gli1 activity (Huang and Schier, 2009). This prompted us to carefully assess the Gli activity distribution along the dorso-ventral axis of the neural tube of *iguana* mutants at the time of lumen remodeling. To that end, we used the TgBAC-(ptch2:Kaede)^a4596^ reporter line (Huang et al., 2012), as ptch2 is a direct target of Gli activity.

By photoconverting Kaede of the TgBAC-(ptch2:Kaede)^a4596^ reporter line at 55hpf in *iguana* and control siblings, the neo-synthetised Kaede pattern at 75hpf reflects the dorso-ventral gradient of Gli activity during the lumen retraction process (Figure 3A) (Jacobs and Huang, 2019). Ventrally, we observed a decrease in the maximal Gli activity in *iguana* compared to controls (Figure 3B, 3B’), as expected in the absence of the primary cilium that is required for maximal Gli activation (Huang and Schier, 2009; Jacobs and Huang, 2019). More dorsally, we observed an increased medial activation of the Gli activity in *iguana* mutant when compared to controls that display a very low level of fluorescence (Figure 3B, 3B’). This strongly suggests that the population of dorsal progenitors that continued to participate abnormally in the lumen border formation in *iguana* has a higher Gli activity than dorsal progenitors in control siblings (Figure 3B, 3B’).

**Figure 3.**
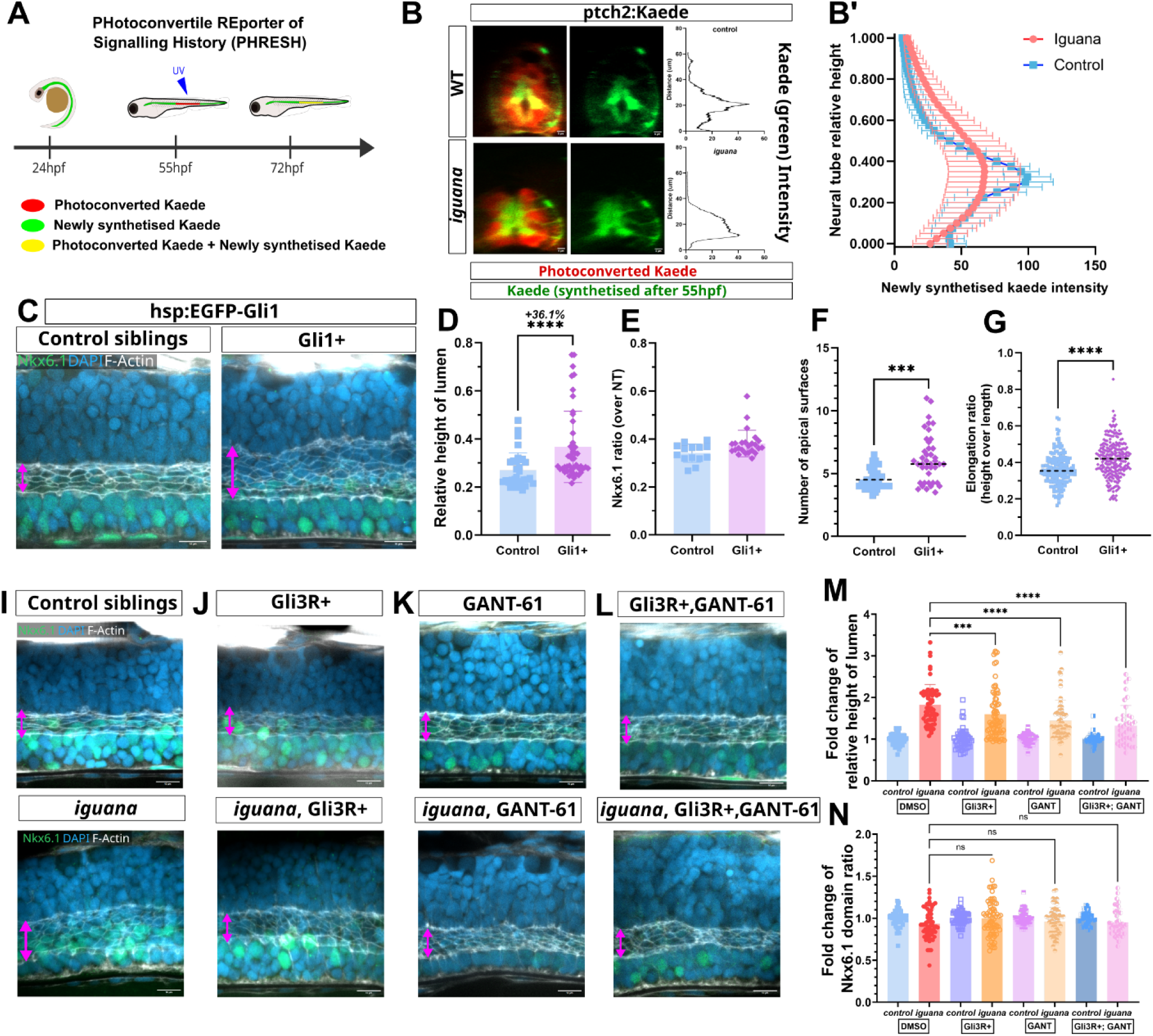
Gli activity is controlling lumen remodelling. **A.** Schematic representation of the experimental design Gli activity is measured using a ptch2:Kaede reporter line. A section of the spinal cord above the yolk extension was photoconverted by the UV light at 55hpf and the fluorescent profile was analysed at 72hpf. We can identify cells that have either new signalling response after photoconversion (green), continued response from before and after photoconversion (yellow), or have ended signalling response before photoconversion (red). **B.** Transverse views of confocal projections of slices of the spinal cord where we photoconverted with an example of graphical signalling profiles from the dorsoventral axis on the green channel, presenting Kaede produce after photoconversion. **B’** Global Hh signalling dynamics by PHRESH analysis. Assessment of Gli activity in control (blue) or iguana (red) embryos along the dorso-ventral axis of the neural tube. **C.** Confocal image of the lumen (F-actin in white) within the neural tube (DAPI in blue) of 75hpf Tg(hsp70l:GFP-gli1)^a4594^ embryos heat-shocked at 40hpf and 50hpf inducing Gli1 over-expression. **D.** Relative lumen height (lumen height over total neural tube height) in controls (blue, n=32) and Gli1 over-expressing embryos (purple, n=46). **E.** Nkx6.1 expression in neural tube (height of Nkx6.1+ domain over total height of the neural tube). **F.** Number of progenitor apical surfaces. **G.** Elongation ratio (height over length) in control siblings (bleu) and Gli1-overexpressed (purple) embryos. **I-J-K-L.** Confocal image of the lumen (white) within the neural tube (blue) in 75hpf Tg(hsp:Gli3R-EGFP)^ca121^ embryos heatshocked at 50hpf to over-express Gli3R (J) or treated at 32hpf with GANT-61 80µM (K) or with both over-expression of Gli3R + GANT-61 treated embryos (L). Pink double arrows represent lumen height. **M.** Graphical representation of lumen size in controls (blue for control n=11 and red for iguana n=10) and Gli3R over-expressing embryos (purple for control n=19 and orange for iguana n=22), GANT-61 treated embryos (light purple for control n=15 and light orange for iguana n=13) and GANT-61 treated embryos (dark blue for control n=19 and pink for iguana n=22). **N.** Graphical representation of Nkx6.1 expression in controls (blue for control n=11 and red for iguana n=10) and Gli3R over-expressing embryos (purple for control n=19 and orange for iguana n=22), GANT-61 treated embryos (light purple for control n=15 and light orange for iguana n=13) and GANT-61 treated embryos (dark blue for control n=19 and pink for iguana n=22). For M and N graphs, each data was normalized by the mean of their relative control condition.

Since the early gradient of Gli activity is known to set the ventral identities of progenitors (Jacobs and Huang, 2019), we wondered if this dorsal shift of Gli activity observed in late development was associated with a dorsal shift of the ventral identities of progenitors. To that end, we immunostained embryos at different stages for Nkx6.1, a marker for ventral progenitor identity (Sander et al., 2000), in *iguana* mutants. When compared to siblings, the higher limit of the Nkx6.1+ domain was only slightly shifted dorsally (Figure S3). Noticeably, the most dorsal progenitors characterised by the absence of Nkx6.1 expression, represented a significant part of the abnormally high lumen border of *iguana* embryos at 72hpf (more than 3 times higher in *iguana* than in control, Figure S3). This indicates that the dorsal cells that persisted in *iguana* dorsal lumen border did not change their identity for Nkx6.1+ ventral progenitors, suggesting the higher lumen height in *iguana* is not the result of simple cell fate change of dorsal to ventral progenitors.

We therefore hypothesised that the absence of Dzip1 in *iguana* mutants leads to abnormally high Gli activity in these dorsal progenitor cells, preventing them from exiting the lumen. Accordingly, we predicted that inducing an ectopic Gli activation at late embryonic stages would prevent lumen retraction. We used a transgenic line over-expressing the activator Gli1 under the control of the heat-shock promoter Tg(hsp70l:GFP-gli1)^a4594^ (Huang et al., 2012). By triggering the induction of Gli1 expression from 40hpf onwards, we observed a defect in lumen retraction : the lumen height at 75hpf of Gli1 expressing embryo was on average 36% higher than heat-shocked controls devoid of Tg(hsp70l:GFP-gli1)^a4594^ (Figure 3C-D). Of note, this late induction of Gli activity did not significantly modify the range of ventral Nkx6.1 expression but did induce a change in the number of progenitor apical surfaces composing the lumen and their apical height-to-length ratio (Figure 3E-G). Hence, Gli1 overactivation before the onset of lumen retraction phenocopies the progenitor behaviour defects of *iguana* mutants. This reinforces our conclusion that, independently of their dorso-ventral identity, a late activation of Gli1 in dorsal progenitors prevents their exit from the lumen and their elongation along the rostro-caudal axis.

We also predicted that reducing the general level of Gli activity in *iguana* mutants should rescue lumen retraction defects. To do so, we first used a heat-shock inducible Gli3R, one of the repressive forms of the Shh pathway, Tg(hsp:Gli3R-EGFP)^ca121^. By triggering the expression of Gli3R just before the onset of lumen remodeling, at 50hpf, we restored a lumen retraction in *iguana* closer to that of the wild-type at 75hpf (Figure 3I-J). Gli3R over-expression from 50hpf onwards had no effect on the wild-type siblings, and did not affect the Nkx6.1 pattern at this late stage where patterning is probably already set (Figure 3M-N, and table S1 for statistical tests). We also submitted *iguana* clutches to pharmacological inhibition of Gli activity with GANT-61, Gli antagonist 61, that interferes with DNA binding of the Gli activator forms (Büttner et al., 2012; Lauth et al., 2007). When treated from 32hpf onwards, the *iguana* mutant lumen was shorter at 75hpf than their counterparts exposed to DMSO (Figure 3K, 3M, and table S1 for statistical tests). Again, this treatment had no effect on lumen border height in wild-type siblings (Figure 3M), nor on Nkx6.1 dorsal border position (Figure 3N, and table S1 for statistical tests). These two experiments present partial rescue of *iguana* phenotype while combining them led to a better rescue of the *iguana* lumen retraction defect (Figure 3L, 3M, 3N, and table S1 for statistical tests). These results indicated that the defects in lumen retraction in *iguana* mutants are caused, at least partially, by an increased Gli activity in dorsal progenitors that form the lumen border. Altogether, our work suggests that the decrease in Gli activity in dorsal progenitors at the onset of spinal remodeling is required for controlling the withdrawal of their apical surface from the lumen, and the shape of the apical surfaces of persisting ventral progenitors.

### 4. The spinal lumen retraction drives roof plate cells extension

Although described in many species, the physical mechanism that coordinates roof plate extension with lumen retraction is not known (Shinozuka and Takada, 2021; Tait et al., 2020). While lumen retracted, the roof plate cells elongated in a coordinated manner. We noticed a slowing down of the roof plate extension concomitant to the defective lumen retraction in *iguana* mutants compared to siblings (GFP signal in green on Figure 2, Figure S2). The coordination between tissues suggests a coupling. To further explore this interaction we sought to accelerate lumen remodeling. Since the lumen border is mostly formed by the apical surfaces of the lateral neural progenitors, we decided to decrease Notch signaling, known to be required for the apical polarisation of neural progenitors from the beginning of lumen formation (Kondrychyn et al., 2013; Sharma et al., 2019). By adding a drug that blocks Notch signaling (LY-411575, a γ-secretase inhibitor) at the onset of lumen remodeling (50hpf), we were able to let the lumen form normally and then determine the effect of a synchronised loss of progenitors apical polarity during lumen remodeling (Figure 4).

**Figure 4.**
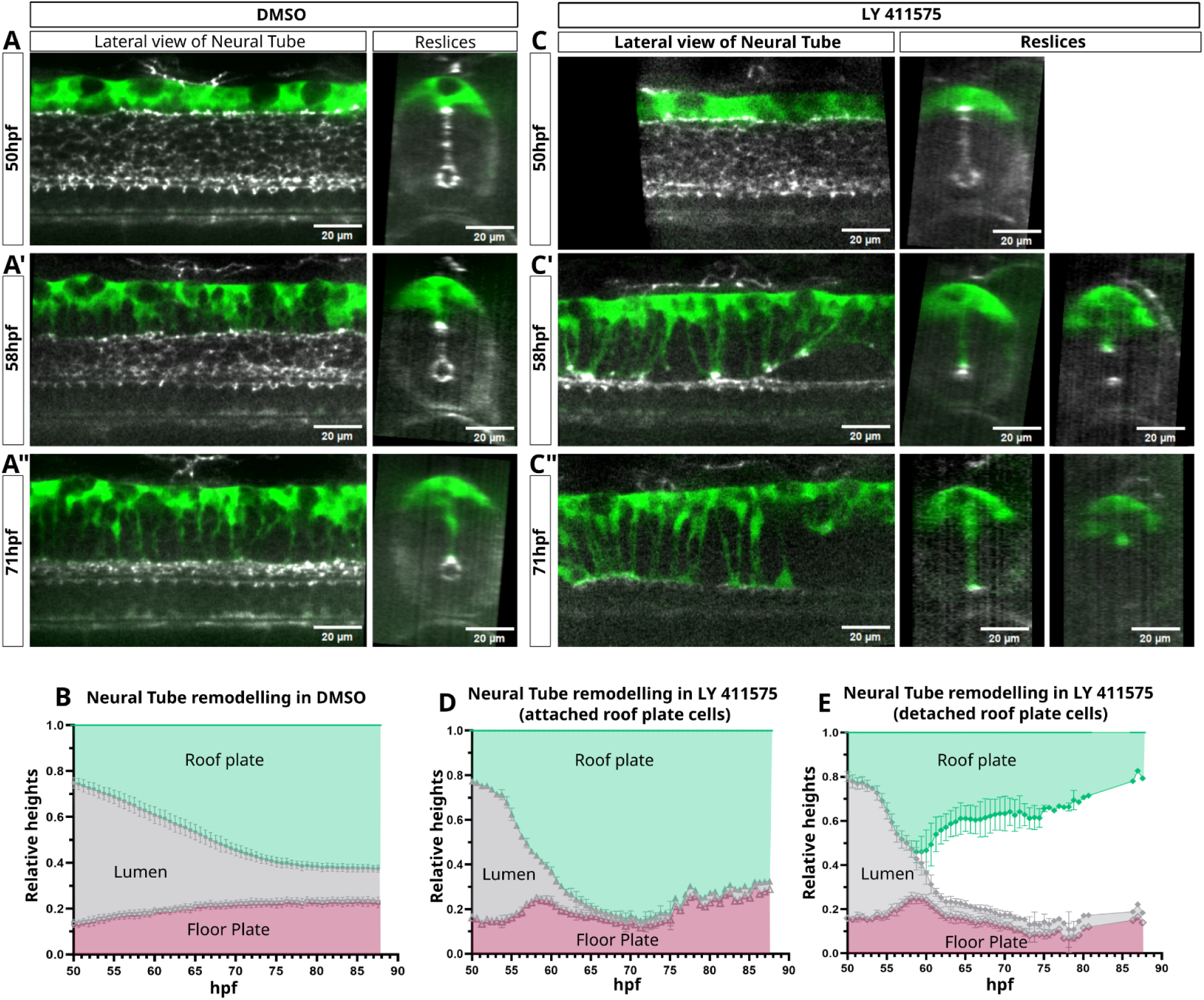
Neural tube remodeling dynamics after treatment inhibiting Notch. Movie images of the neural tube of transgenic control embryos Et(krt4:eGFP)^sqEt33^, TgKi(tjp1a-tdTomato)^pd1224^ treated with 0.05% DMSO (A, A‘, A’) and embryos treated with LY-411575 (25µM), a Notch inhibitor (C, C‘, C”) at 50hpf, 60hpf and 75hpf, lateral and transverse views. Two distinct behaviours are observed in the RP cell population, with some cells detaching from the lumen and others remaining attached to the lumen (C‘’). Graphical representation of the movie with analysis of the size of the different tissues making up the neural tube, RP cells in green, lumen in grey, FP cells in purple in control embryos (B, n = 10 embryos) or embryos treated with LY 411575 (D, E, n = 6 embryos). Separate analyses of the two RP cell behaviours, RP cells that remain attached to the lumen (D) and RP cells that detach from the lumen (E). Lumen borders retract faster in LY condition and RP cells elongate at the speed of lumen border retraction. We can observe a slight deformation of the FP around 57hpf, when RP cells start to detach from the lumen border.

Long term real-time confocal imaging allowed us to describe the whole remodeling process in DMSO-treated control embryos, starting from a fully extended lumen border dorso-ventrally and a small dorsal roof plate (50hpf, Figure 4A). We observed that the roof plate cells extended dynamic processes concomitantly to lumen retraction, with their endfeet continuously forming a dorsal ceiling of the lumen along the rostro-caudal axis, consistently with previous observations (Kondrychyn et al., 2013 and Figure S2). This remodeling stopped around 75hpf, when the central canal persisted ventrally (Figure 4A’’). As described previously, by measuring the dorso-ventral height of the different tissues, we were able to assess a remodeling speed of lumen roughly 1µm/h between 50 and 75hpf (Figure 2B, Figure S2B, Figure 4B see Figure S5A and Material and methods for details). We then compared with embryos treated with LY-411575 from the beginning of our time-lapse imaging (49hpf – Figure 4C), and we observed as anticipated an acceleration of lumen retraction, including the loss of the most ventral apical surfaces that normally persist (compare Figure 4A’ and C’). We evaluated that the lumen retraction was more than three times faster (from ∼1.25 µm/h in DMSO-treated to ∼4 µm/h in LY-treated, Figure 4B and 4D, Figure S5A). The roof plate cell extension presented the same acceleration of lumen retraction and proceeded faster than normal for the first 10 hours of filming (Figure 4C’, D-E). We then observed two types of behaviour : (i) the majority (66%, 203 cells out of 307, in 13 embryos) of roof plate processes retracted dorsally (Figure 4C’’, 4D), following a loss of contact with the apical material of the fastly retracting lumen border, giving rise to a middle spinal space without a lumen (polarised progenitor cells were mostly replaced by extra differentiated neurons as a result of Notch inhibition; Figure S5E’-G). (ii) Some roof plate processes (33%) kept a constant attachment to the apical surfaces of the fastly retracting lumen, eventually forming a contact with the floor plate cells that appeared to be resistant to Notch signal loss (Figure 4C”, 4E, S5E”). In both cases, we observed a decrease of Sox2+ cells number and an increase of HuC+ cells number meaning a loss of progenitors due to LY-411575 treatment (Figure S4F-G).

We reasoned that the retraction following the loss of contact with apical surfaces forming the lumen might be the result of the roof plate processes “springing back” after losing the force that extended them. Indeed, we noticed that the floor plate behaviour was modified upon LY-411575 treatment. In the wild-type controls, the floor plate cells exhibited a minimal change of height compared to the roof plate cells (Figure 4A-B). However, upon acceleration of the lumen retraction by Notch inhibition, the floor plate cells were abnormally stretched, which could reflect an increased traction force exerted by the fastly retracting lumen (compare movie 2, and Figure 4 B, D and E for quantification). Consistently, the floor plate cells also changed their height in movement resembling a relaxation when the majority of roof plate cells detached from the lumen around 59hpf (Figure S4D-E for quantification). Altogether, our results argue that the lumen retraction is generating the forces that drive roof plate cells extension through a mechanical coupling between the two tissues.

## DISCUSSION

In this work, we showed that Gli activity regulation is central for the retraction of the spinal lumen and that this retraction is the driving force of roof plate cell elongation. It illustrates the importance of fine-tuned coordination between cell behaviours in the control of morphogenetic processes involving different cell types to achieve the harmonious development of a tissue.

### Cilia are not required for lumen border remodeling, despite their importance in spinal development

We observed that the lumen border remodeling proceeded normally without cilia, even in ciliary mutants that had an impact on lumen inflation (Figure 1 and S1) (Thouvenin et al., 2020). This suggests that the size of the inflated ventral part of the lumen where the CSF circulates does not control the final height of the central canal that results from lumen remodeling. Our results with the *foxj1a* mutant failed to reproduce the morpholino phenotype targeting *foxj1a* mRNA, which displayed a transitory delay in lumen retraction (Ribeiro et al., 2017), maybe due to technical differences in our respective approaches. Our results with *foxj1a* mutants are consistent with the other mutants we used to deplete motile (*dnaaf1*) or all (*elipsa*) cilia, comforting us in our conclusion that neither motile nor primary cilia are directly required for lumen remodeling.

We noted that *dnaaf1* mutants displayed an abnormal elongation of the floor plate domain inducing a slight dorsal shift of the central canal (Figure 1F’), as it has been observed in *foxj1a* morphants in Ribeiro et al. 2017. This suggests that motile cilia are not necessary to control lumen border retraction but might be implicated in the proper position of the canal dorso-ventrally by controlling either the elongation of floor plate cells or the differentiation of neurons deriving from ventral progenitors.

### Dzip1 acts upstream Gli activity in lumen retraction control

The *iguana* mutant is an exception among the ciliary mutants we observed, as its lumen border failed to fully retract (Figure 1). While manipulating *iguana* embryos, we noticed a minor but reproducible deformation of their whole neural tube. The *iguana* mutants presented a neural tube with a slightly shorter dorso-ventral axis while slightly larger in medio-lateral axis (Figure S1C-E-E’). We think that this small neural tube deformation is the consequence of a general change in the trunk mechanical properties, as it was more prominent when we mounted live embryos in agarose for live imaging compared to fixed samples. This could be a consequence of the role of cilia in somite development (Huang and Schier, 2009). Several pieces of argument prompted us to decouple this general change of embryonic shape from the specific lumen retraction defects we studied : (1) The neural tube deformation precedes a significant lumen border height difference between *iguana* and controls (Figure S1, before 50hpf). (2) The lumen border in *iguana* is higher than in control when measured in absolute height (µm on Figure S2), the opposite of what would be expected in a shorter neural tube. (3) We also observed a slightly shorter neural tube in the *elipsa* mutant, but no lumen retraction defect was observed (Figure 1 and S1). (4) When we mimicked the *iguana* phenotype with an overactivation of Gli, we observed a change of lumen border height independently of an effect on neural height (Figure 3C).

Our PHRESH experiment in the *iguana* spinal cord results suggest a change in the dorso-ventral gradient of Gli at the time of lumen remodeling (Figure 3B-B’), where a low-level Gli activity is progressing more dorsally in the absence of Dzip1. The rescue experiments showing that reducing Gli activity restores a shorter lumen in *iguana* mutants confirms that the phenotype we observed is a consequence of an abnormal activity of Gli preventing lumen border retraction. Our results therefore demonstrate a role of Dzip1 in regulating the Gli activity gradient for controlling spinal lumen remodeling.

### Dzip1 regulates the Gli activity gradient at the onset of lumen remodeling

Dzip1 is required for the early maintenance of primary cilia, which are known to be crucial to control the post-translational modifications of the Gli proteins to control the balance between activator and inhibitor forms (Andreu-Cervera et al., 2021). In the zebrafish embryo muscles and neural tube, it has been suggested that the early absence of cilia prevents the maximal Gli activation, while promoting a larger domain of low Gli activity, via the transcriptional regulation of Gli1 (Huang and Schier, 2009). This dorsal extension of low activity might in part rely on the absence of Gli repression dorsally, as cilia are also essential for the processing of Gli repressor forms (Andreu-Cervera et al., 2021). Our PHRESH experiment (Figure 3A-C) indicates that this persists between 2 and 3 days post fertilization (dpf) when the lumen remodeling takes place. However, the absence of lumen retraction defects in the *elipsa* mutant indicates a specific requirement for Dzip1 in regulating Gli.

This difference might be due to a specific role of Dzip1 in zebrafish ciliogenesis. The major difference of the *iguana* mutant compared to other ciliary mutants is the earlier loss of primary cilia in the *iguana* mutant due to a minimal maternal contribution, as supported by previous works and ours (Figure S1) (Omori et al., 2008; D’Gama et al., 2024). One possibility is that the loss of primary cilia at early stages of development triggers an aberrant Gli gradient regulation that later persists until the stage of lumen remodeling, due to the ability of Gli1 to act on its own promoter (Jacob and Briscoe, 2003), therefore perpetuating a defect caused by early loss of cilia in iguana.

Alternatively, we cannot exclude that Dzip1 is required in setting the Gli activity gradient independently of its role in ciliogenesis. One report described a dual role of Dzip1 in regulating the Gli activity in HEK293, including via direct interaction between Dzip1 and Gli3, preventing this repressor form of Gli to localise to the nucleus (Wang et al., 2013). Regardless of the mechanism, our results show that Dzip1 specifically controls Gli gradient activity in late developmental stages to control the spinal lumen remodeling.

### A new role for the spinal Gli activity gradient in regulating morphogenesis

While Gli activity was already known to regulate proliferation and patterning of the spinal progenitor cells (Sagner and Briscoe, 2019), our work proposes that the regulation of Gli activity is also essential for lumen retraction control. This control appears largely independent of the early role of Gli in setting the upper limit of Nkx6.1 expression, suggesting that most of our manipulation of the Gli gradient did not significantly affect patterning the differentiation of neural progenitor cells.

It was recently reported that the extent of the Gli activity domain in the zebrafish embryo spinal cord decreases ventrally from 55hpf to 72hpf, a period that coincides with the lumen remodeling (Jacobs and Huang, 2019). In addition, Canizares et al. (2020) observed in mice neural tube a decline of Shh activity concomitant with central canal retraction. Our different experiments in the zebrafish *iguana* mutant and with the ectopic expression of Gli activator all show that an excess of Gli activity prevents lumen retraction. We therefore propose that the gradual decrease of Gli activity in the most dorsal progenitors is a prerequisite for their disengagement of the lumen border, independently of a change of cell fate. Interestingly, this suggests that progenitor proliferation, patterning and cell behaviour are under the control of the same gradient, allowing synchronised cell cycle exit, differentiation and morphogenesis downstream of the same Gli effectors.

### Mechanisms downstream of Gli controlling lumen retraction

Shh signaling has already been suggested to control neural tube morphogenesis in the past, namely by regulating the apical domain size in the ventral midline of the folding hindbrain in mice (Brooks et al., 2020, 2025). Interestingly, the apical area of progenitors forming the lumen in zebrafish is not affected when we manipulate Gli activity, but its height-to-length ratio is (Figure 2), arguing for different mechanisms downstream of Shh signaling in these different systems. Although slightly different, these two mechanical changes regulated by Gli activity are commonly dependent on a contractile acto-myosin network (Lecuit and Le Goff, 2007). It suggests that effectors downstream of Gli signaling can regulate the mechanical properties of this acto-myosin network of epithelial progenitor cells.

The precise mechanism for the disengagement of dorsal progenitors from the lumen border, as well as the downstream effectors of Gli controlling this behaviour, are not known and will be the subject of further studies. Noticeably, when we experimentally decreased Gli activity in *iguana* mutants to rescue their lumen border phenotype, we never observed the extrusion of ventral progenitor that form the final central canal, as it is the case when we inhibited Notch signaling (compare Figure 3J-K-L and Figure 4). This suggests that dorsal and ventral progenitors regulate the engagement of their apical domain into the lumen border differently. It is consistent with the observation in different species that the final central canal is formed by a majority of Nkx6.1 positive progenitors, that correspond to the most ventral progenitor (Figure S3) (Cañizares et al., 2020; Fu et al., 2003; Ribeiro et al., 2017; Tait et al., 2020). The signaling pathway(s) affected by Gli activity in the control of progenitor engagement in the lumen are not known. One hypothesis is that it converges with Notch signaling at some level, as Notch is a very important signal for maintaining apical polarisation of spinal progenitors (Sharma et al., 2019), and Shh and Notch are interacting at different levels by cross-regulation (Jacobs and Huang, 2021).

During early neurogenesis, it has been shown that polarised radial glia cells that engage into neural (or oligodendrocyte lineage) differentiation extrude from the lumen, sometimes by severing their apical domain, as shown in chicken embryos (Das and Storey, 2014; Saade and Martí, 2025). However, self-renewal of polarised progenitors maintains the extent of the lumen along the dorso-ventral axis during most of the neurogenesis period. One can hypothesise that the same extrusion behaviour is recapitulated at the end of neurogenesis by terminating dorsal progenitors and contributes to lumen border size reduction.

To explain this lumen reduction, a previous study suggested that during neurogenesis, the most ventral progenitors can disengage from the lumen through terminal differentiation and that more dorsal progenitors replenish the ventral population of polarized progenitors (Ravanelli and Appel, 2015). On the opposite, another study has proposed a model whereby the most dorsal radial glia cells delaminate in an ordered manner at the end of neurogenesis, giving rise to the lumen retraction that resembles a ratchet (Tait et al., 2020). A more stochastic process can also be envisioned. To date, our results do not allow us to decide the precise position and order of the disengagement of polarized progenitors in the remodeling zebrafish spinal lumen. The dynamics of this cell behaviour leading to spinal lumen retraction in our zebrafish model will require further studies, to fully characterise the link between individual cell extrusion and global lumen retraction.

In the literature, the microRNA miR-219 has been involved in the control of the size of the lumen. When blocked by a morpholino, the lumen retraction failed (Hudish et al., 2016, 2013). This miR-219 is targeting a series of genes encoding proteins required for apical polarisation, like Prcki and Pard3, suggesting that its expression is required at the end of neurogenesis for progenitors withdrawal from the lumen border (Hudish et al., 2013). These works identified Shh signaling as a regulator of lumen retraction (Hudish et al., 2016), consistently with our own results. However, all the manipulation of the Shh were performed from early developmental stages, which suggests that they might change the whole patterning of the spinal cord progenitor. In our case, we manipulated Gli activity later in development, and always monitored that the ventral limit of Nkx6.1 - a marker for ventral progenitors - was not massively shifted dorsally (Figure S3). Therefore, the precise link between miR-219, apical domain regulation and Gli activity needs to be reexamined in the future. Nevertheless, together with these previous works, our results lead us to propose that the decrease of the Gli activity in the dorsal part of the spinal cord is required to promote the dorsal progenitors behaviour associated with apical domain disengagement from the lumen border, leading to its retraction.

### Retraction of the lumen is the driving mechanical force leading to roof plate cells extension

As mentioned in introduction, the concomitance of the roof plate cell extension with lumen retraction was already well described in the past (Böhme, 1988; Cañizares et al., 2020; Kondrychyn et al., 2013; Ribeiro et al., 2017; Ševc et al., 2009; Shinozuka and Takada, 2021; Tait et al., 2020). However, the driving force of this remodeling process remained an open question (Shinozuka and Takada, 2021; Tait et al., 2020).

By manipulating Notch signaling at the beginning of lumen remodeling, we managed to accelerate lumen border retraction, and showed that the roof plate cells were accelerating their extension. This argues for the lumen border retraction being the pulling force onto the roof plate cells, which are prone to deformation in response to this stress. Our conclusions are comforted by a series of observations : (1) previous reports showed that the roof plate (and floor plate) cells do not lose their radial glia identity when Notch was blocked (Kondrychyn et al., 2013; Sharma et al., 2019); it is therefore unlikely that our LY-411575 treatment modified the roof plate cells behaviour directly. This is further confirmed by the fact that the lumen retraction speed upon LY-411575 treatment is constant, independently of whether roof plate cells remained attached to the lumen or recoiled back (Figure 4). We therefore conclude that Notch inhibition primarily affects lumen border retraction, and that the roof plate responds to this increased pulling force via mechanical coupling. (2) This hypothesis is further reinforced by the behaviour of roof plate cells that failed to keep their attachment to the lumen border. At this stage, we have no evidence that can explain why some cells behaved this way, whereas others managed to keep a constant attachment to the lumen border. However, once detached, these roof plate cells retracted their extension in a way that resembles a spring recoiling back, arguing for the fact that they experienced tension from the lumen retraction. This could be further examined in the future with the help biophysical modeling. (3) Although floor plate cells seemed resistant to large deformations, we noticed that they slightly increased their height in response to lumen retraction acceleration, arguing for a pulling force being applied symmetrically on both extremities of the lumen. This slight deformation is also released when a large fraction of roof plate cells recoiled, as if this detachment of roof plate cells was liberating the excessive force applied by the retracting lumen onto the floor plate. The molecular nature of the mechanical attachment between the roof plate and the lumen, and the biophysical properties allowing roof plate cells to elongate cellular extension will be the scope of further studies.

It has been proposed that, at least in chicken and mouse, roof plate cells secrete a soluble version of the Crumbs2 protein that promotes apical disengagement of the dorsal progenitors (Tait et al., 2020). This is compatible with our observations in the zebrafish embryo, in a model that would suggest that a secretion from the roof plate participates in the lumen retraction that in return, exerts a pulling force on the roof plate cells to trigger their elongation.

Altogether, our work offers new evidence of the importance of mechanical coupling in a morphogenesis process involving several cell types. Interestingly, the zebrafish spinal cord remodeling allowed us to describe quantitatively the effect of this mechanical coupling together with refining our understanding of the signaling regulation of neural progenitor behaviours by Gli. Our system opens new avenues to describe and model the detailed control of morphogenesis that is required for harmonious spinal cord development in vertebrates.

## ACKNOWLEDGMENTS

We want to thank Okan Ayas, François Ozon et Matthieu Antoine-Le Louët for their help while doing short internships in the lab. We are grateful to Daniel Levic and Michel Bagnat for sharing published fish lines, and EZRC for sending the Et(krt4:eGFP)^sqEt33^ line. The Dextran for injection was kindly provided by Marie Breau. We thank Christine Vesque, Marion Baraban and Marie Breau for their fruitful advice on the manuscript.

This work was supported by the following institutions: Sorbonne Université (Emergences SU 2019) ; Groupement des Entreprises Françaises dans la lutte contre le Cancer (GEFLUC 1356) ; Ligue Contre le Cancer régionale (Ile de France) ; Fondation pour la Recherche Médicale (équipe FRM : EQU202503020010) ; Association pour la Recherche sur le Cancer (ARC - 4th year of PhD). Natural Sciences and Engineering Research Council of Canada (RGPIN-2022-03167).

## Movies Legendes

**Movie 1. Lumen remodelling in a typical control embryo.** This movie presents images of a live time-lapse confocal acquisition from 50hpf until 85hpf of the spinal neural tube of a non-mutant transgenic embryo : Et(krt4:eGFP)^sqEt33^;TgKi(tjp1a-tdTomato)^pd1224^ . Roof plate cells are shown in green by cytoplasmic EGFP inserted in the sqEt33 locus and the apical surfaces of progenitor cells are outlined in gray by a fusion of the junctional protein ZO-1 fused to tdTomato. A stack of images was acquired every 15 minutes, average projections are shown.

**Movie 2. Lumen remodelling in a typical iguana embryo.** This movie presents images of a live time-lapse confocal acquisition from 50hpf until 85hpf of the spinal neural tube of an iguana mutant transgenic embryo : Et(krt4:eGFP)^sqEt33^;TgKi(tjp1a-tdTomato)^pd1224^ . Roof plate cells are shown in green by cytoplasmic EGFP inserted in the sqEt33 locus and the apical surfaces of progenitor cells are outlined in gray by a fusion of the junctional protein ZO-1 fused to tdTomato. A stack of images was acquired every 15 minutes, average projections are shown. Notice the higher lumen at the end of the process and the slower extension of roof plate cells.

**Movie 3. Lumen remodelling in a typical control embryo treated with DMSO.** This movie presents images of a live time-lapse confocal acquisition from 50hpf until 75hpf of the spinal neural tube of a transgenic embryo treated with 0.05% DMSO : Et(krt4:eGFP)^sqEt33^;TgKi(tjp1a-tdTomato)^pd1224^ . Roof plate cells are shown in green by cytoplasmic EGFP inserted in the sqEt33 locus and the apical surfaces of progenitor cells are outlined in gray by a fusion of the junctional protein ZO-1 fused to tdTomato. A stack of images was acquired every 15 minutes, average projections are shown.

**Movie 4. Lumen remodelling in a typical control embryo treated with LY-411575.** This movie presents images of a live time-lapse confocal acquisition from 50hpf until 75hpf of the spinal neural tube of a transgenic embryo treated with LY-411575 at 25µM : Et(krt4:eGFP)^sqEt33^;TgKi(tjp1a-tdTomato)^pd1224^ . Roof plate cells are shown in green by cytoplasmic EGFP inserted in the sqEt33 locus and the apical surfaces of progenitor cells are outlined in gray by a fusion of the junctional protein ZO-1 fused to tdTomato. A stack of images was acquired every 15 minutes, average projections are shown. Notice the faster retraction of the lumen, the concomitant acceleration of roof plate cells elongation and the loss of contact of some roof plate cells with the lumen border accompanied by a dorsal retraction of their extension.

---

**Figure S1.**
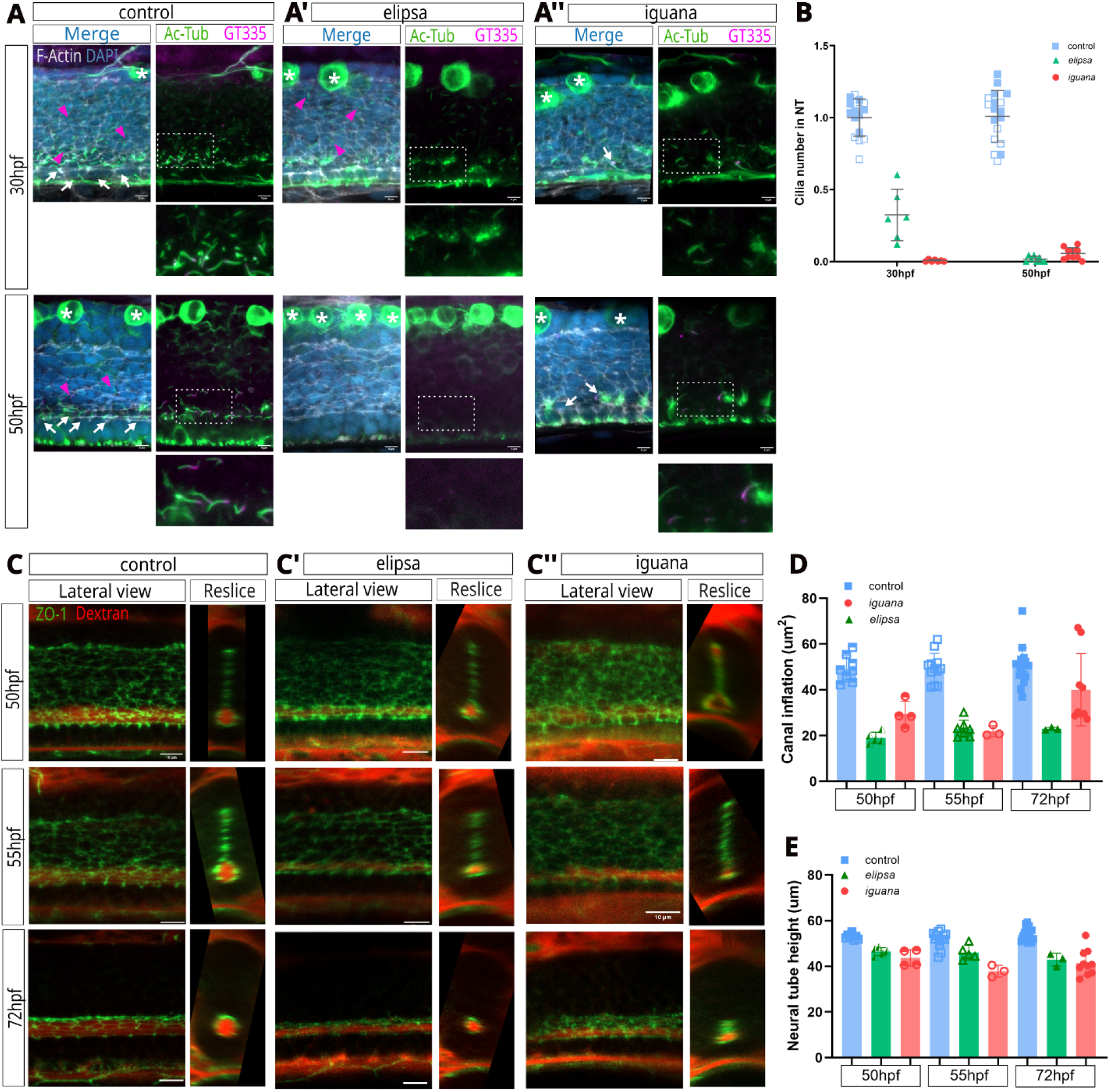
Comparison of phenotypes between ciliary mutants iguana and elipsa. **A.** Confocal images of immunostainings for cilia axonemes (ac-tub for acetylated tubulin – green) and basal bodies (GT335 for glutamylated-tubulin, magenta), shown on lateral views of the spinal cord recognized by its dense nuclear DAPI staining (blue). Note that the large dorsal Rohon Beard neurons have a strong cytoplasmic ac-tub signal independent of cilia (asterisk). In control siblings at 30 and 50hpf (n=16 and n=17), we observed a widespread distribution of short primary cilia along the dorso-ventral axis (pink arrowheads on left panels), and a specific ventral enrichment in longer cilia that correspond to the motile cilia (white arrows on left panels) (Borovina et al., 2010). **B.** Quantification of the relative cilia density on the different mutant conditions at 30 and 50hpf - quantifications are plotted in reference to the control number of cilia. We observe a 75 % decrease in the cilia density in elipsa at 30hpf, probably reflecting a maternal contribution of elipsa mRNA (n=6, pink arrowhead on upper middle panel) (Omori et al., 2008). On the contrary, the iguana cilia density is close to zero with only a few ventral motile cilia persist (n=7, white arrow in top right panel), consistent with previous results (Huang and Schier, 2009). At 50hpf, at the onset of lumen remodeling, elipsa mutants were completely devoid of cilia (n=8, lower middle panel) while in iguana we could detect no primary cilia and a few ventral motile cilia, sometimes together with aberrant bundles of tubulin-based protrusion (n=10, white arrow in lower right panel). **C.** Confocal imaging of live embryos where the apical sides of progenitor cells forming the lumen is labelled by ZO1-GFP (green) and the circulation cavity of the lumen is filled with a TexasRed-Dextran (red - see (Thouvenin et al., 2020). The Dextran also fills extracellular spaces around the neural tube. Typical lateral views of spinal cords at 50, 55 and 75hpf are shown, together with transverse views reconstructed from the confocal stacks. These images were used to measure neural tube height, canal height and canal inflation. **D-D’.** Canal inflation measurement in mutants and control siblings. On the transverse images of the spinal cord at the three time points, the diameter of the lumen circulating canal was measured and represented as bar plots for the elipsa mutants (D - green, n=5 at 50hpf, n=5 at 55hpf, n=3 at 72hpf) or iguana mutants (D’ - red, n=4 at 50hpf, n=3 at 55hpf, n=10 at 72hpf) and their siblings (in blue, n=8 at 50hpf, n=14 at 55hpf, n=17 at 72hpf). Consistent with previous works showing the role of motile cilia in lumen inflation (Thouvenin et al., 2020) we observed a decrease in the diameter of the two ciliary mutants at all stages. **E-E’.** We also measured and plotted the total height of the neural tube in the same genetic conditions and time points, and we noted a small consistent decrease of the neural tube height in the two mutants elipsa (E) and iguana (E’). We therefore chose to normalise most of our height measurement in ciliary mutants by the total height of the neural tube to compensate for these small variations.

**Figure S2.**
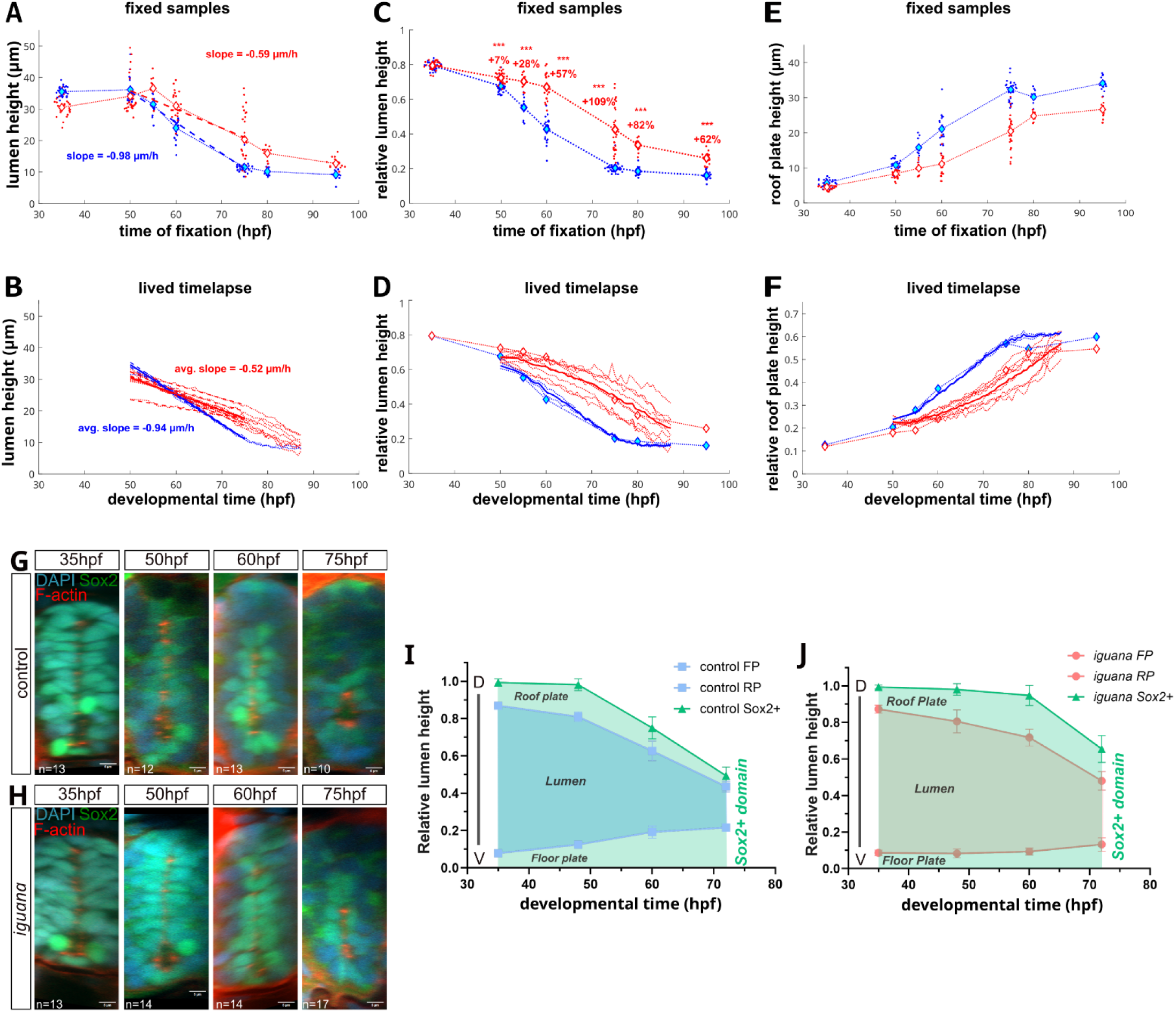
Quantification of lumen height lived and fixed and Sox2+ cells in iguana. **A.** Raw data in µm of lumen height along time in fixed control (blue) and iguana (red) embryos. Linear fits between 55 and 75hpf are shown as dashed lines, their slope representing the speed of retraction. **B.** Raw data live timelapse in µm of lumen height over time, in control (blue) and iguana (red) embryos. Linear fits between 55 and 75hpf are shown as bold lines, their slope representing the speed of retraction. **C.** Relative lumen height (lumen height over total neural tube height) in iguana (red) and control siblings (blue).The percentage of increase in the iguana mutant is indicated above each timepoint. **D.** Relative lumen height of live timelapse and fixed embryos with corrected time for live, iguana (red) and control siblings (blue). **E.** Raw data in µm of roof plate height along time in fixed control (blue) and iguana (red) embryos. **F.** Relative roof plate height (roof plate height over total neural tube height) in iguana (red) and control siblings (blue).The percentage of increase in the iguana mutant is indicated above each timepoint. Large dots represent the mean of fixed data and the dashed line the average of each genotype, while dotted lines are lumen retraction of each embryo in the live experiment. The mean of the control and embryos lumen retraction is in bold line. A good agreement between fixed and live experiments is found. A-F For fixed embryos, at 35 hpf, n=28 controls and n= 25 iguana ; at 50hpf, n=25 controls and n= 26 iguana; at 55 n=11controls and n= 10 iguana ; at 60 hpf, n=22 controls and n= 22 iguana ; at 75 hpf n= 30 controls and n= 27 iguana : at 80 hpf, n=12 controls and n= 11 iguana ; at 95 hpf, n=15 controls and n= 16 iguana. For live timelapse movies, n=2 controls and n=8 iguana. **G-H.** Sox2 expression in control **(G)** and iguana **(H)** neural tube at 35hpf, 50hpf, 60hpf, 75hpf. Sox2 expression is in green, F-actin (stained with phalloidin, red) and nucleus (stained with DAPI, blue). **I-J.** Quantification of Sox2 expression domain height along the neural tube dorso-ventral axis in control **(I)** and iguana **(J)**. Graphs represent each part of the neural tube (floor plate, lumen, roof plate) within their relative height to the neural tube. Lumen height is highlighted in blue (control) or red (iguana) and the Sox2+ domain is highlighted in green along the dorso-ventral axis. We can observe a persistence of progenitors Sox2+ cells in iguana in the dorsal region.

**Figure S3.**
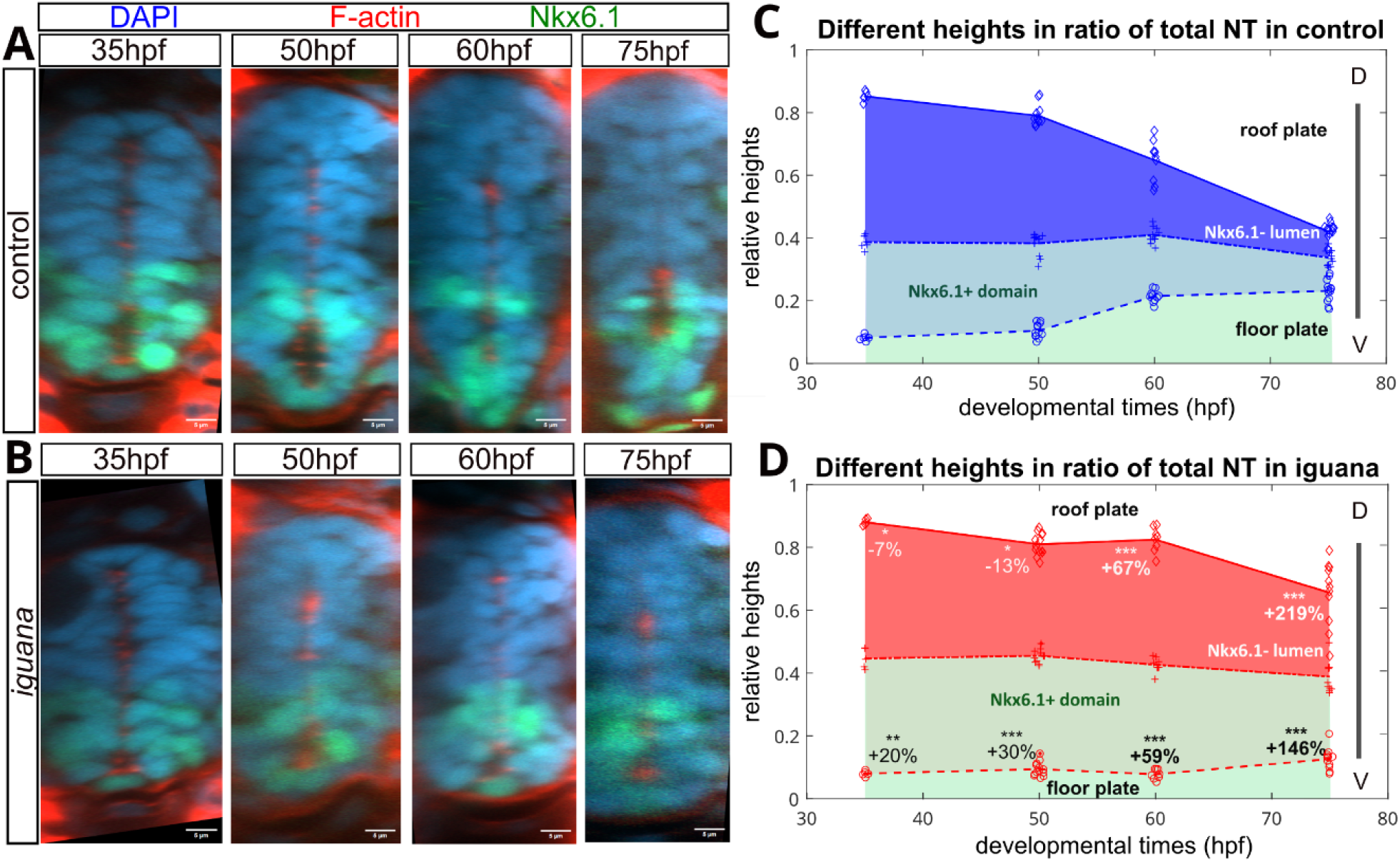
Nkx6.1 expression in control and iguana neural tube. **A-B.** Confocal images of neural tube (at the level of the cloaca) at 35hpf, 50hpf, 60hpf, 75hpf. Nkx6.1 expression is in green, F-actin (stained with phalloidin) is in red and nuclei (stained with DAPI) are in blue. **C-D.** Quantification of the Nkx6.1 expression domain along the neural tube dorso-ventral axis in control (n=6 at 35hpf, n=12 at 50hpf, n=10 at 60hpf, n=13 at 75hpf) and iguana (n=5 at 35hpf, n=13 at 50hpf, n=8 at 60hpf, n=11 at 75hpf). Graphs represent each part of the neural tube (floor plate, lumen, roof plate) within their relative height to the neural tube. We highlight the lumen part and distinguish the Nkx6.1- domain of the lumen with dark color and Nkx6.1+ domain of the lumen in light color. Percentage represents the difference between control and iguana at each stage for either the Nkx6.1- or Nkx6.1+ domain. See (Sander et al., 2000)

**Figure S4.**
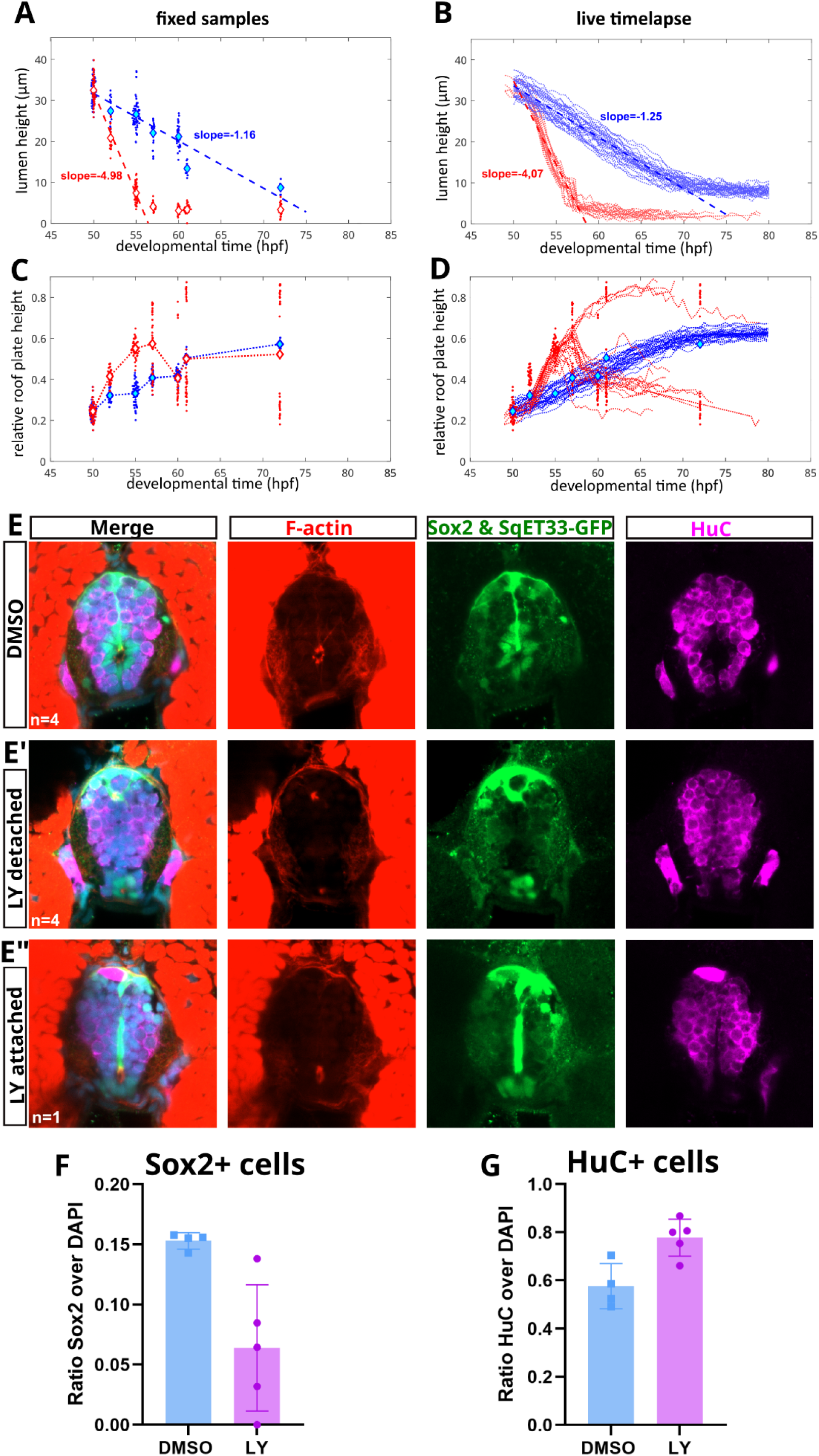
LY411575 induces progenitor differentiation. **A.** Lumen height in µm, measured in fixed embryos treated by DMSO (blue) or LY-411575 (25µM, red) from 50hpf to 72hpf. **B.** Lumen height in µm, measured in live embryos treated by DMSO (blue) or LY-411575 (25µM, red) from 50hpf to 80hpf, one image is taken every 15 min. **C.** Relative height of roof plate in the neural tube, measured in fixed embryos treated by DMSO (blue) or LY-411575 (25 µM, red) from 50hpf to 72hpf. **D.** Relative height of roof plate in the neural tube, measured in live embryos treated by DMSO (blue) or LY-411575 (25 µM, red) from 50hpf to 80hpf, one image is taken every 15min. A-D. Fixed embryos : 50 hpf, n= 55 slices of DMSO and 55 slices of LY-411575 ; 55 hpf, n= 21 slices of DMSO and 30 slices of LY-411575 ; 55 hpf, n= 62 slices of DMSO and 63 slices of LY-411575 ; 57 hpf, n= 21 slices of DMSO and 32 slices of LY-411575 ; 60 hpf, n= 33 slices of DMSO and 25 slices of LY-411575 ; 61 hpf, n= 12 slices of DMSO and 43 slices of LY-411575 ; 72 hpf, n= 9 slices of DMSO and 31 slices of LY-411575. Live imaging : n=19 slices in 6 embryos from DMSO, n=30 in 10 embryos in LY-411575. **E-E’-E”.** Confocal images of neural tube of 72hpf Et(krt4:eGFP)^sqEt33^ zebrafish embryos in DMSO (E, n=4) or treated with LY411575 (25 µM, n=5) from 50hpf to 72hpf (E’-E”). Roof plate cells are GFP+ in green in the dorsal part of the neural tube and the extension is along the spinal midline. The central canal is stained with phalloidin (F-actin, red), progenitor cells are Sox2 (green) and differentiated cells HuC (far red). In LY411575 embryos, two distinct behaviours are observed in the RP cell population, some RP are detaching from the lumen (E’) and others remaining attached to the lumen (E”). **F.** Ratio of Sox2+ cells over DAPI+ cells in the neural tube of DMSO and LY-411575 embryos. **G.** Ratio of HuC+ cells over DAPI cells in DMSO and LY-411575 conditions. In both cases, we observed a reduced number of Sox2+ cells (F) and an increase of HuC+ cells (G).

**Table S1.** Tables of statistics for Tg(hsp:Gli3R-EGFP)^ca121^ treated with GANT-61 experiment, a two-way ANOVA were performed with multiple comparisons. Statistics are done with GraphPad (Prism). Mean difference, summary of significance and adjusted p-value of each comparison is shown in those tables. On the left, the table of lumen height comparison between each condition of the experiment and on the right the Nkx6.1 domain analysis. ns = p>0.05; * = p<0.05; ** = p<0.01; *** = p<0.001; **** = p<0.0001.

| Table1 |  |  |  |
| --- | --- | --- | --- |
| Lumen Height comparison | Mean diff. | Summary | Adjusted P Value |
| control DMSO vs. iguana DMSO | -0,1677 | **** | <0,0001 |
| control DMSO vs. control HS-GFP | 0,009004 | ns | 0,9984 |
| control DMSO vs. control GANT | -0,01597 | ns | 0,9577 |
| control DMSO vs. control GANT HS-GFP | 0,01302 | ns | 0,9951 |
| iguana DMSO vs. iguana HS-GFP | 0,05812 | ** | 0,0012 |
| iguana DMSO vs. iguana GANT | 0,04272 | ns | 0,068 |
| iguana DMSO vs. iguana GANT HS-GFP | 0,1197 | **** | <0,0001 |
| control HS-GFP vs. iguana HS-GFP | -0,1186 | **** | <0,0001 |
| control HS-GFP vs. control GANT | -0,02498 | ns | 0,7471 |
| control HS-GFP vs. control GANT HS-GFP | 0,004012 | ns | >0,9999 |
| iguana HS-GFP vs. iguana GANT | -0,01539 | ns | 0,9756 |
| iguana HS-GFP vs. iguana GANT HS-GFP | 0,06163 | ** | 0,0054 |
| control GANT vs. iguana GANT | -0,109 | **** | <0,0001 |
| control GANT vs. control GANT HS-GFP | 0,02899 | ns | 0,6855 |
| iguana GANT vs. iguana GANT HS-GFP | 0,07702 | *** | 0,0002 |
| control GANT HS-GFP vs. iguana GANT HS-GFP | -0,06098 | ** | 0,0089 |
| Nkx6.1 domain comparison | Mean diff. | Summary | Adjusted P Value |
| control DMSO vs. iguana DMSO | 0,02311 | ns | 0,1448 |
| control DMSO vs. control HS-GFP | 0,006577 | ns | 0,9971 |
| control DMSO vs. control GANT | -0,01683 | ns | 0,6631 |
| control DMSO vs. control GANT HS-GFP | -0,02344 | ns | 0,4539 |
| iguana DMSO vs. iguana HS-GFP | -0,004275 | ns | 0,9998 |
| iguana DMSO vs. iguana GANT | -0,02746 | ns | 0,086 |
| iguana DMSO vs. iguana GANT HS-GFP | -0,02786 | ns | 0,2107 |
| control HS-GFP vs. iguana HS-GFP | 0,01226 | ns | 0,8757 |
| control HS-GFP vs. control GANT | -0,02341 | ns | 0,3163 |
| control HS-GFP vs. control GANT HS-GFP | -0,03002 | ns | 0,1561 |
| iguana HS-GFP vs. iguana GANT | -0,02318 | ns | 0,3302 |
| iguana HS-GFP vs. iguana GANT HS-GFP | -0,02359 | ns | 0,4043 |
| control GANT vs. iguana GANT | 0,01248 | ns | 0,8888 |
| control GANT vs. control GANT HS-GFP | -0,006613 | ns | 0,999 |
| iguana GANT vs. iguana GANT HS-GFP | -0,0004044 | ns | >0,9999 |
| control GANT HS-GFP vs. iguana GANT HS-GFP | 0,01869 | ns | 0,719 |

